# 40S ribosomal subunits move along the 5ʹ UTR through an eIF4A-independent mechanism in higher eukaryotes

**DOI:** 10.64898/2026.06.23.734070

**Authors:** Thomas Ossevoort, Olga Chernaya, Angela Browning, Dave H. Mathews, Dmitri N. Ermolenko

## Abstract

During translation initiation, the 40S small ribosomal subunit is recruited to the mRNA 5′ cap and scans the 5′ untranslated region (UTR) to locate the start codon. While the mechanism of 40S translocation remains elusive, the RNA helicase eIF4A has long been suspected as the primary molecular motor driving 40S scanning. In this study, we utilized GFP reporter mRNAs to investigate the impact of 5′ UTR length on translational efficiency. We found that an 8-fold variation in the length of unstructured 5′ UTRs did not lead to substantial changes in translation efficiency in wheat germ extract (WGE) and human HEK293T cell lysate. By contrast, the presence of a stable stem-loop in the middle of the 5′ UTR significantly reduced cap-dependent translation. These results suggest that mRNA scanning is not rate-limiting when the UTR is devoid of secondary structure. Inhibition of eIF4A by hippuristanol in cell-free protein synthesis systems yielded an equivalent decrease in translation for mRNAs with short and long unstructured 5′ UTRs, indicating that eIF4A may be dispensable for 40S scanning. Our data suggest that helicase-independent one-dimensional diffusion may be the primary mechanism enabling 40S movement along the 5′ UTR during initiation.

## Introduction

The initiation of eukaryotic translation is a multi-step process that serves as a primary regulatory checkpoint for cellular gene expression. Translation initiation begins with the formation of the eIF4F complex, a heterotrimer consisting of the cap-binding protein eIF4E, the scaffolding protein eIF4G, and the DEAD-box RNA helicase eIF4A^1^. The eIF4F complex facilitates mRNA recruitment by recognizing the 5′ 7-methyl guanosine (m^7^G) cap. The 43S pre-initiation complex (PIC), which consists of the 40S ribosomal subunit, various initiation factors, and the eIF2-GTP-Met-tRNAi ternary complex^2^, is then recruited by the eIF4F complex to the 5′ end of mRNA to form the 48S PIC.

Following mRNA recruitment, the assembled 48S PIC moves along the 5′ untranslated region (5′ UTR) to locate the first AUG codon downstream of the 5′ end via a process called scanning^3–6^. Start site recognition triggers the release of initiation factors, the 60S ribosomal subunit joining and formation of the 80S ribosomal complex ^7–10^.

Despite its central role in eukaryotic initiation, the mechanism governing 48S PIC movement through the 5′ untranslated region (5′ UTR) remains poorly defined. Whether ribosomal scanning relies on directional ATP-dependent translocation or is entirely governed by one-dimensional diffusion remains a fundamental and unresolved question^11^. It is commonly assumed that eIF4A, which is the only ATPase and helicase among the core initiation factors, serves as the directional motor for 40S translocation along the 5′ UTR^12–16^. However, *in vitro* studies have demonstrated that scanning does not depend on ATP concentration^14^ and can occur in an eIF4A-independent manner^17,18^.

During translation initiation, the 40S ribosomal subunit must navigate a diverse landscape of 5′ UTRs with varying lengths and complex secondary structures. Despite the inherent versatility of the translation machinery, 5′ UTR lengths are notably constrained across eukaryotes when compared to 3’ UTRs. While 3’ UTRs frequently span over a thousand nucleotides, 5′ UTRs are significantly shorter. For example, median lengths of 5′ UTRs in budding yeasts and humans are ∼50 and 220 nucleotides, respectively^19^. The evolutionary ceiling on 5′ UTR length may indicate that scanning processivity is a limiting factor for translation initiation.

Multiple studies have investigated the dependence of scanning time and/or translational efficiency on the 5′ UTR length to interrogate the mechanism of scanning^14,20–24^. Some reports revealed a negative correlation between 5’ UTR length and protein synthesis^22,23,25^. Others showed that translation efficiency is independent of 5’ UTR length^22,24^ or is even enhanced by lengthening of the 5’ UTR^20,21^. Many of these studies utilized natural 5′ UTRs and thus are confounded by the inherent propensity of RNA to adopt secondary structures, which impede 48S scanning complexes independent of 5′ UTR length^20,21,23–27^.

To decouple the effects of 5′ UTR length from the influence of RNA secondary structure, unstructured 5′ UTRs, such as CAA repeats, can be employed^28,29^. However, the high propensity of CA and CAA repeats to recombine and their low stability during the propagation of mRNA-encoding plasmids *in vivo* restricts the range of 5′ UTR lengths that can be explored. We have recently developed an algorithm for designing <u>n</u>on-repetitive <u>u</u>nstructured <u>s</u>equences (NUS) that are significantly longer than repetitive unstructured 5′ UTRs used in previous studies^30,31^. We found that extending the unstructured 5′ UTR of GFP reporters 10-fold over the median length of yeast 5′ UTRs reduces GFP synthesis in yeast cells by less than two-fold^31^, indicating that 40S scanning is not rate-limiting in lower eukaryotes. Here, we employ the GFP reporter mRNAs with unstructured, nonrepetitive 5′ UTRs to investigate whether 40S scanning constrains the rate of protein synthesis in higher eukaryotes. We demonstrate that translation efficiency is largely independent of 5′ UTR length in human and plant cell extracts. Furthermore, inhibition of the main translational helicase eIF4A equally reduces translation of mRNAs with long and short unstructured 5′ UTRs, indicating that eIF4A is dispensable for 40S scanning. Our data support the model suggesting that 40S progression along the 5′ UTR is governed predominantly by one-dimensional diffusion rather than ATP-driven translocation.

## Results

### Translational efficiency in wheat germ extract is independent of 5′ UTR length

To test the impact of 5′ UTR length on translational efficiency in higher eukaryotes, we utilized a series of superfolder GFP (sfGFP) reporters containing non-repetitive unstructured sequences (NUS) in the 5′ UTR as previously described (Figure 1A)^31^. The NUS1 5′ UTR sequences (^50^NUS1-^400^NUS1), ranging from 50 to 400 nucleotides (nts), were computationally engineered to lack secondary structure while maintaining substantial sequence complexity. The NUS1 5′ UTR sequences were also devoid of guanosines, except for the first two Gs added at the 5′ end for effective *in vitro* transcription. In addition, UCC and ACCAC sequence motifs, which were previously shown to inhibit translation in yeast^32^ were avoided. A series of 5′ UTR variants were generated by progressively truncating the 3′ end of the original 400 nt NUS sequence while preserving the original Kozak context of the start codon (Fig. 1A). The GFP ORF was followed by the 3′ UTR from yeast *RPL41B* mRNA and a 30-nucleotide-long poly(A) tail. GFP reporter mRNAs were transcribed by T7 run-off transcription and enzymatically capped with 5′ m^7^Gppp cap.

**Figure 1.**
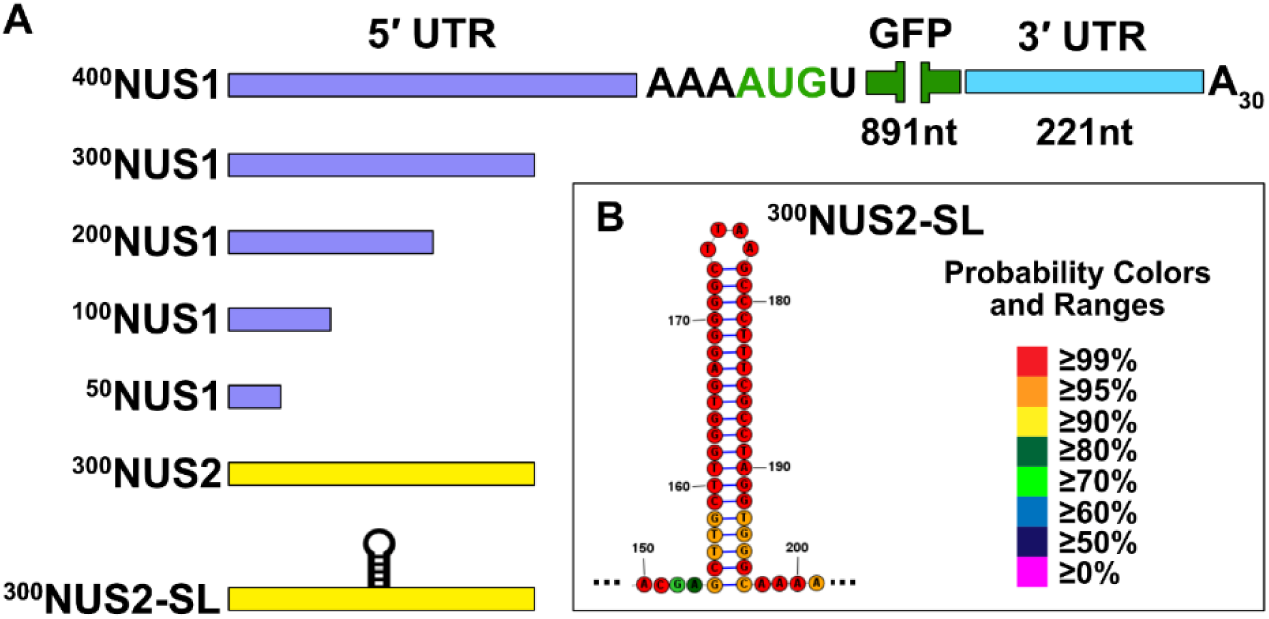
Design of GFP reporter mRNAs containing unstructured (NUS) 5′ UTRs. A) The NUS1 5′ UTRs (purple) share a common 5′ cap proximal sequence and range in length from 50 to 400 nt. The unstructured 300 nt-long ^300^NUS2 (yellow) has a distinct sequence composition. ^300^NUS2-SL features a stable stem-loop (shown in panel B) inserted into the middle of the 300 nt-long 5′ UTR. All constructs contain a common Kozak consensus sequence flanking the start codon of sfGFP ORF, the 3’ UTR derived from yeast RPL41B mRNA and a 30 nt-long poly(A) tail. The base-pairing probability of the stem-loop (B) of ^300^NUS2-SL determined by the RNAstructure software is shown by color code as indicated.

To evaluate the translational efficiency of our unstructured 5′ UTR GFP reporters, we utilized the commercially available plant cell-free protein synthesis system, wheat germ extract (WGE). WGE, which is widely used to study protein synthesis, is a robust, highly active and cap-dependent translation system^33^. By utilizing WGE, we can decouple protein synthesis from confounding processes such as transcription, splicing, and mRNA export, thereby isolating translational output as a direct function of 40S scanning efficiency.

To examine cap dependence of WGE, we compared the translational efficiency of the ^300^NUS1 sfGFP reporter with and without a 5′ m^7^Gppp cap (Figure 2A). After a ∼10-minute lag period, which at least in part is due to GFP folding and chromophore maturation^34–36^, GFP fluorescence increased linearly until its rise slowed down at ∼150-180 minutes. The well-known autoinhibition of cell extracts is due to depletion of components essential for protein synthesis, such as ATP, GTP and amino acids. The relative translational efficiency of capped and uncapped mRNA was quantified by comparing the slopes of the linear increase of fluorescence intensity between 30 and 120 minutes. The absence of the 5′ cap resulted in >12-fold reduction in sfGFP fluorescence (Figure 2B, Suppl. Table 1). The linear accumulation of sfGFP signal indicates a constant translation rate and thus the lack of mRNA decay^37^. Our previous studies have also demonstrated that transcripts remain stable and do not undergo significant degradation within 2 hours of translation in WGE^38^. Consistent with these observations, levels of both capped and uncapped mRNAs determined by RT-qPCR remained unaffected after 120 minutes of translation in WGE (Suppl. Figure 1A). Hence, an order of magnitude reduction in GFP synthesis from uncapped mRNA shows that translation in WGE is predominantly dependent on the presence of 7-methyl-guanosine mRNA cap.

**Figure 2.**
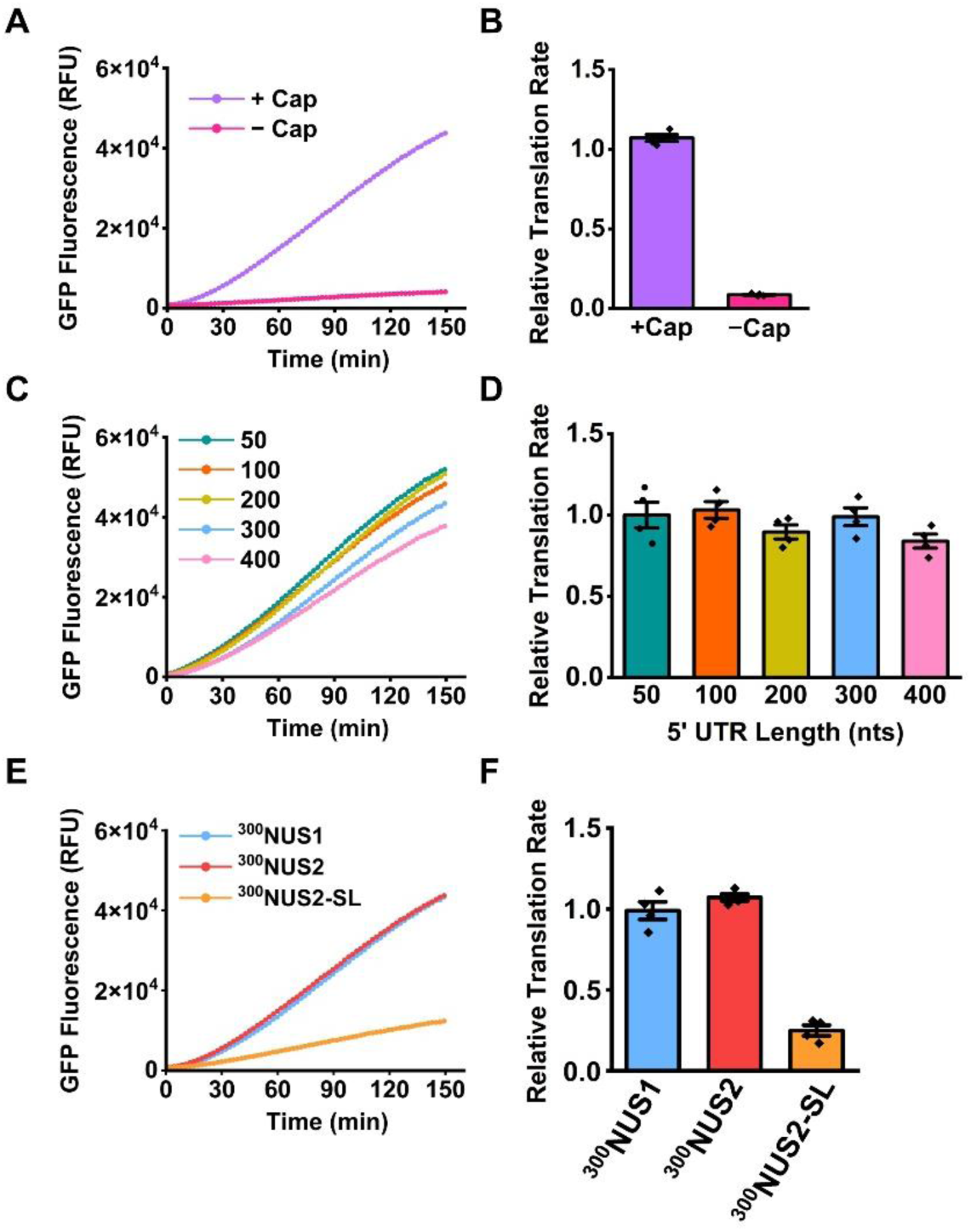
40S scanning in WGE is independent of 5′ UTR length and sequence. (A, C, E) GFP fluorescence measured (relative fluorescent units, RFU) as a function of time during reporter mRNA translation in WGE. (B, D, F) Relative translation rates determined by the slopes of linear fluorescence growth in the range of 30-120 minutes and normalized to the slope observed with ^50^NUS1 mRNA. Bar graphs represent mean values; error bars show the standard error of the mean (SEM) calculated from three to four independent translation experiments. P-values indicating statistical significance of observed changes in translational efficiency are shown in Supplemental Table 1. (A-B) Translation of capped and uncapped ^300^NUS1 mRNAs. (B-C) Translation of ^50–400^NUS1 mRNAs. (D-E) Translation of ^300^NUS1,^300^NUS2 mRNA and ^300^NUS2-SL mRNAs.

To assess the impact of 5′ UTR length on translational efficiency, we measured translation of ^50^NUS1-^400^NUS1 mRNA reporters with unstructured 5′ UTRs spanning 50 to 400 nts (Figure 2C-D). It is noteworthy that the 400 nt-long 5′ UTR is ∼twice as long as the mean length of 5′ UTRs in wheat (235 nt)^39^ and three to four times longer than the median 5′ UTR lengths reported for other plant species, such as tomato (106 nt) or maize (132 nt)^19^. Slopes of the linear increase of fluorescence intensity between 30 and 120 minutes were normalized to the slope observed for ^50^NUS1 mRNA to determine the relative translation rate of each reporter construct (Figure 2D). Remarkably, an 8-fold increase in 5′ UTR length from 50 to 400 nts had no significant impact on translation efficiency (Figure 2D, Suppl. Table 1). This data suggests that 5′ UTR length is not rate-limiting for translation initiation in WGE.

To investigate how sequence variations in the unstructured 5′ UTR affect GFP synthesis, we examined translation of ^300^NUS2 mRNA with a 300-nt-long unstructured 5′ UTR, which was designed independently from ^300^NUS1 as previously described^31^. ^300^NUS1 and ^300^NUS2 shared as much sequence identity as two shuffled sequences^31^. There was no significant difference in GFP synthesis from ^300^NUS1 and ^300^NUS2 reporters, indicating that variations in unstructured sequences did not affect translation efficiency (Figure 2E-F, Suppl. Table 1).

We next explored how the presence of secondary structure, which is thought to impede 40S scanning^20,21,23–27^, affects translation of reporter mRNAs. Nucleotides 156-199 of ^300^NUS2 5′ UTR were replaced with a GC-rich stem-loop composed of 20 base pairs and a UAAA tetraloop to create ^300^NUS2-SL reporter mRNA as previously described^31^. This stem-loop is expected to hamper 40S scanning without affecting 40S recruitment because this element of secondary structure is positioned over 150 nt downstream from the 5′ end of mRNA, i.e. much further away than the ∼45 nt footprint of the 43S initiation complex^40,41^. Insertion of the stem-loop resulted in a 4-fold decrease in GFP synthesis (Figure 2E-F). RT-qPCR measurement revealed no significant degradation of ^300^NUS1, ^300^NUS2 and ^300^NUS2-SL mRNAs, indicating that stem-loop insertion reduced translation rather than mRNA stability (Suppl. Figure 1B). This result indicates that our GFP reporters are translated in a scanning-dependent manner rather than by internal initiation (e.g., IRES-mediated initiation).

### Translational efficiency in human cell extract is independent of 5′ UTR length

To determine whether the relationship between 5′ UTR length and translational output is conserved in humans, we extended our analysis to human HEK293T cell-free translation system^42^. To prevent eIF2α phosphorylation, which inhibits translation initiation and is often induced in cell cultures during lysate preparation, we used genetically modified HEK293T cells that express GADD34, a eIF2α-specific regulatory subunit of human PP1 phosphatase, and vaccinia virus K3L protein, a substrate for eIF2α-specific kinases, as previously described^42^. To verify the cap-dependency of the HEK293T lysate system, we compared the translational efficiency of the ^300^NUS1 reporter with and without a 5′ m^7^Gppp cap (Figure 3A). In the absence of m^7^Gppp cap, GFP levels were undetectable by fluorescence (Figure 3B). RT-qPCR shows that one-hour incubation of uncapped mRNA in HEK293T lysate leads to a ^~^25-fold decrease in mRNA levels, while the capped mRNA remained intact (Suppl. Figure 2A). Hence, mRNA translation and stability in HEK293T cell extract strictly requires the presence m^7^Gppp cap.

**Figure 3.**
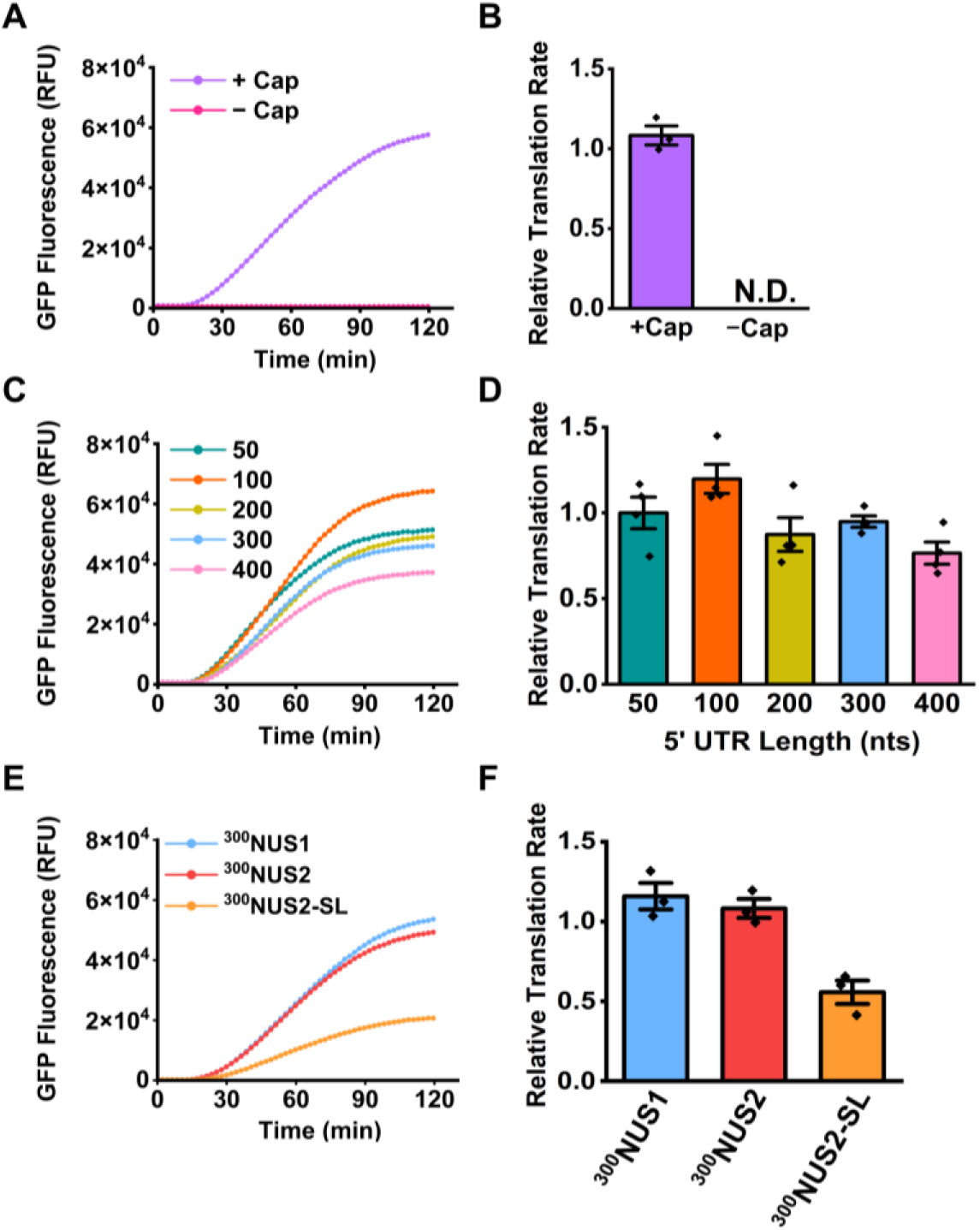
Translation efficiency in HEK293T human cell lysates is insensitive to changes in the length of the unstructured 5′ UTR. (A, C, E) GFP fluorescence measured (relative fluorescent units, RFU) as a function of time during reporter mRNA translation in HEK293T lysate. (B, D, F) Relative translation rates determined by the slopes of linear fluorescence growth in the range of 20-60 minutes and normalized to the slope observed with ^50^NUS1 mRNA. Bar graphs represent mean values; error bars show the standard error of the mean (SEM) calculated from three to four independent translation experiments. p-values indicating statistical significance of observed changes in translational efficiency are shown in Supplemental Table 2. (A-B) Translation of capped and uncapped ^300^NUS1 mRNAs. (B-C) Translation of ^50–400^NUS1 mRNAs. (D-E) Translation of ^300^NUS1,^300^NUS2 mRNA and ^300^NUS2-SL mRNAs.

To assess the impact of 5′ UTR length on translational efficiency, we measured the output of mRNA reporters featuring unstructured 5′ UTRs spanning 50 to 400 nucleotides (Figure 3C). The translational output of each sfGFP reporter was quantified by calculating the slope of the linear phase of fluorescence from 20 to 60 minutes and normalized to the ^50^NUS1 sfGFP construct to calculate relative translation efficiency. Similar to WGE, an 8-fold increase in 5′ UTR length had a minimal impact on total protein synthesis (Figure 3C-D, Suppl. Table 2). A very modest (<20%), statistically significant reduction of GFP synthesis was only observed with the ^400^NUS1 mRNA as compared to the ^50^NUS1 mRNA (Figure 3D, Suppl. Table 2). ^300^NUS1 and ^300^NUS2 mRNAs were translated with similar efficiencies, demonstrating that variations in the sequence of unstructured 5′ UTR had little impact on protein synthesis in HEK293T cell extract (Figure 3E-F). By contrast, a stable 20-bp stem-loop in the middle of the 300 nt-long 5′ UTR reduced translation two-fold (Fig. 3E-F). RT-qPCR showed that the levels of GFP reporter mRNAs remained unchanged during translation in HEK293T lysates (Suppl. Figure 2B-C).

Taken together, our experiments in WGE and HEK293T extract show that 40S scanning along is 5′ UTR not-rate limiting for protein synthesis unless 40S movement is impeded by mRNA secondary structure.

### eIF4A helicase activity is dispensable for 40S scanning through unstructured 5′ UTRs

The DEAD-box RNA helicase eIF4A is thought to be the primary ATP-dependent helicase necessary for eukaryotic translation initiation. While it is established that eIF4A facilitates the loading of the 43S pre-initiation complex to most cellular transcripts^29,41,43,44^, it remains unclear whether eIF4A also drives 40S scanning. To evaluate the role of eIF4A in 40S scanning, we utilized hippuristanol an allosteric eIF4A inhibitor to test whether mRNAs with longer 5′ UTRs are more sensitive to eIF4A inhibition because of eIF4A involvement in scanning. Hippuristanol targets the C-terminal domain of eIF4A, effectively inactivating the helicase by blocking the conformational rearrangement coupled to ATP hydrolysis^45,46^.

Because hippuristanol IC_50_ was reported to be highly transcript and cell-type dependent^37,46^, we monitored GFP synthesis programmed by ^50^NUS1 mRNA across a range of hippuristanol concentrations (Suppl. Figure 3A). Based on these results, we selected 0.5 µM and 1 µM concentrations, at which hippuristanol inhibits GFP expression by ∼ 50% and 80%, respectively. We did not find a significant difference in the reduction of GFP synthesis between ^50^NUS1, ^200^NUS1 and ^400^NUS1 mRNAs at either 0.5 µM or 1 µM hippuristanol concentration (Figure 4A-B, Suppl. Table 3). Corresponding RT-qPCR measurements showed no significant changes in mRNA levels during translation in the presence of hippuristanol (Suppl. Figure 3B), indicating that hippuristanol inhibits protein synthesis without affecting mRNA stability in WGE. Hence, while eIF4A activity is required for protein synthesis, it is dispensable for the 40S scanning during translation initiation.

**Figure 4.**
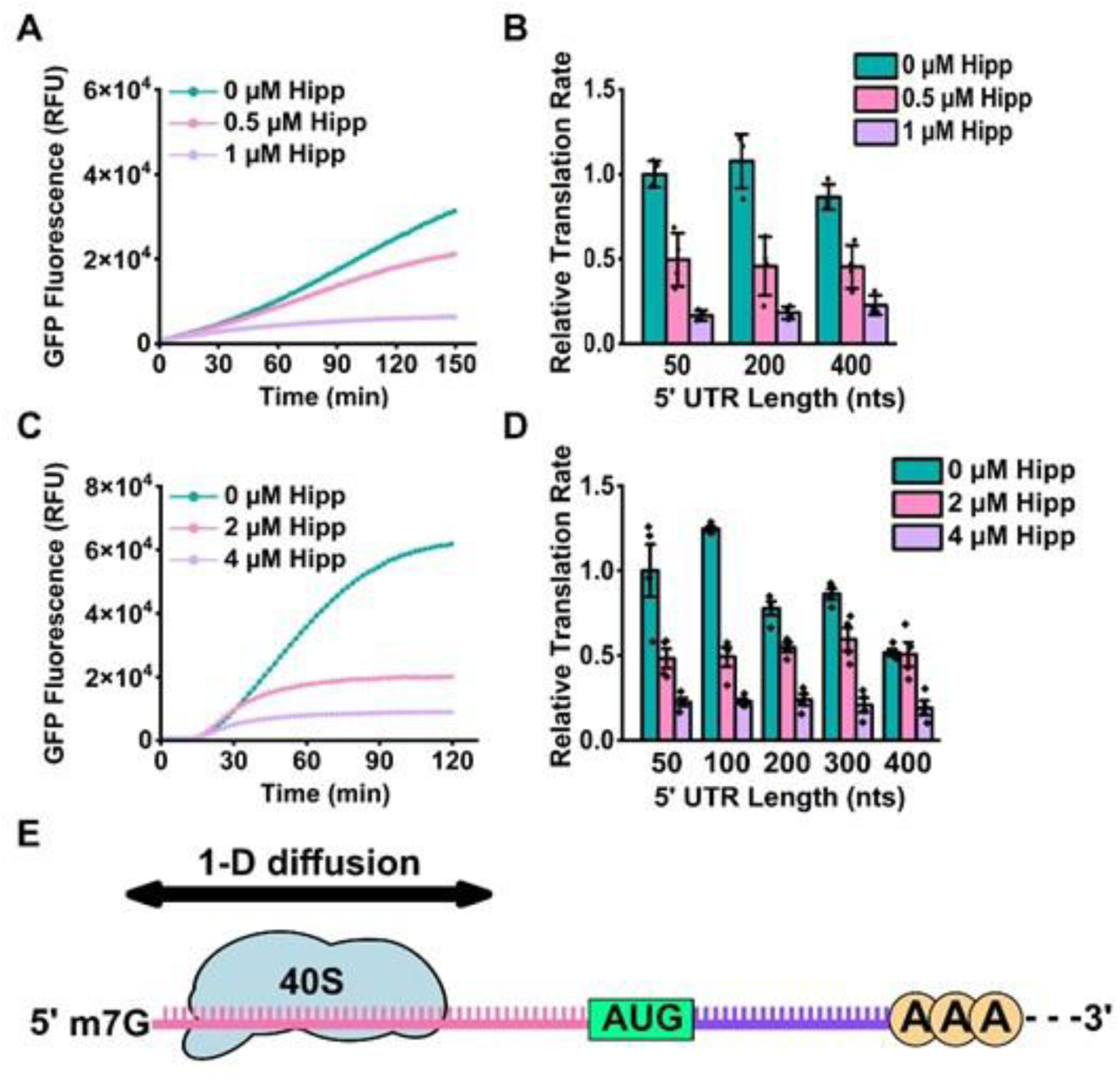
eIF4A inhibition by hippuristanol uniformly reduces translation of reporter mRNAs with short and long 5′ UTRs in WGE and HEK293T lysate. (A, C) GFP fluorescence measured (relative fluorescent units, RFU) as a function of time during ^50^NUS1 mRNA translation in WGE (A) or in HEK293T cell lysate (C) in the presence of 1% DMSO and several concentrations of hippuristanol as indicated. (B, D) Relative translation rates determined by the slopes of linear fluorescence growth and normalized to the slope observed with ^50^NUS1 mRNA in the presence of 1% DMSO and absence of hippuristanol. The length of NUS1 5′ UTR is shown on the x-axis. Bar graphs represent mean values; error bars show the standard error of the mean (SEM) calculated from three to four independent translation experiments. p-values indicating statistical significance of observed changes in translational efficiency are shown in Supplemental Tables 3 (WGE) and 4 (HEK293T). (E) Schematics depicting 40S scanning of the 5′ UTR in search of the start codon by 1D diffusion.

Next, we measured inhibition of ^50^NUS1 mRNA translation by various concentrations of hippuristanol in HEK293T lysate (Suppl. Figure 3C) and found that 2 µM and 4 µM hippuristanol inhibited GFP synthesis by roughly 50% and 80%, respectively. Similar to our observations in WGE, treatment of HEK293T lysates with 2 µM and 4 µM hippuristanol resulted in a uniform decrease of GFP expression programmed by ^50^NUS1, ^200^NUS1 and ^400^NUS1 mRNAs, regardless of 5′ UTR length (Figure 4C-D). RT-qPCR data indicated that 2 µM and 4 µM hippuristanol treatment in HEK293T lysate induced a 2-fold and 4-fold decrease in GFP mRNA levels, respectively (Suppl. Figure 3D, Suppl. Table 4). The observed decrease in mRNA levels is likely due to the coupling of downregulation of translation initiation to mRNA decay. Indeed, cap-binding eIF4F complex (eIF4E•eIF4G•eIF4A) is known to compete for binding with decapping machinery and protect mRNA from decapping and 5′ to 3′ exonucleolytic decay^47–49^. Nevertheless, observed in both WGE and HEK293T cell extracts, equal reduction of GFP expression from mRNAs with short and long 5′ UTRs induced by hippuristanol suggests that eIF4A is dispensable for 40S scanning.

## Discussion

In this work, we investigated the mechanistic basis of evolutionary constraints on the 5′ UTR length and possible involvement of eIF4A in 40S scanning. By utilizing GFP reporters with unstructured 5′ UTRs, we demonstrate that an 8-fold increase in 5′ UTR length from 50 to 400 nts has a modest effect on translation, if any, in both plant and human-derived translationally active cell lysates. Notably, a 400 nt-long 5′ UTR is nearly two-fold longer than the median length of 5′ UTRs in plants and humans^19^. Our data are consistent with experiments that manipulated the length of unstructured 5′ UTRs in yeast cells^24,31^, and indicate that mRNA scanning by the 40S subunit is not a rate-limiting step of translation initiation in both lower and higher eukaryotes.

In contrast to the lengthening of the unstructured 5′ UTR, cap-distal secondary structure in the 5′ UTR was observed in our and other studies^24,31,50–53^ to strongly inhibit translation. Hence, the negative correlation between 5′ UTR length and protein synthesis observed in previous studies^23,25^ and the evolutionary constraints on 5′ UTR length^19^ are likely due to the increased extent of secondary structure in longer 5′ UTRs that hinders 40S scanning^31,54^.

Our data also suggests that 40S scanning is unlikely to be driven by eIF4A helicase activity, as translation of GFP reporters with short and long 5′ UTRs are equally reduced by the eIF4A inhibitor, hippuristanol. eIF4A is critical for the eIF4E•eIF4G•eIF4A binding to the 5′ cap and the following recruitment of the 43S PIC onto the mRNA^17,29,44,55,56^. However, eIF4A was shown to be dispensable for 40S scanning along unstructured 5′ UTRs in both this and previous studies^17,31,57^. Other helicases, such as DDX3X and DHX29, are needed for translation on mRNAs with highly structured 5′ UTRs^12^. However, recapitulation of translation initiation and 40S scanning in systems reconstituted from purified components, which do not include additional helicases besides eIF4A^14,17^, indicates that 40S movement along the 5′ UTR is a helicase-independent, diffusion-based process.

One-dimensional (1D) diffusion is a common mechanism by which DNA and RNA-binding proteins search for their target sites^56,58^. The ability of the 40S to migrate along mRNA in a helicase-independent manner suggests that 1D diffusion of the 40S is not restricted to unstructured 5′ UTRs used in this and other reports^17,31,57^, but also occurs on natural mRNAs. Consistent with the 1D diffusion model of 40S scanning, the bi-directional movement of the 40S subunit along the 5′ UTR was supported by several *in vivo* studies^59–61^ and observed in single-molecule *in vitro* experiments^14^. Based on our data and a large body of published observations reviewed in detail elsewhere^31^, we thus conclude that 1D diffusion is a parsimonious explanation of 40S movement along the 5′ UTR during mRNA scanning.

## Materials & Methods

### mRNA variant Design and Plasmid Construction

DNA sequences encoding the various sfGFP reporters, which were originally described elsewhere^31^, were subcloned into the modified pSP64 Poly(A) vector (Promega) downstream of a T7 promoter. Non-repetitive unstructured sequences (NUS) were designed using the orega (Optimizing RNA Ends with a Genetic Algorithm) tool within the RNAstructure software package (https://rna.urmc.rochester.edu/RNAstructure.html)^30^. The algorithm iteratively evolved sequences using a scoring function that penalizes base-pair formation while maximizing linguistic complexity. For sequences still predicted to contain base-pairs after the initial run, calculations were restarted for additional evolution. Base-pair probabilities were verified using the partition function in RNAstructure. All reporter mRNA sequences are shown in the Suppl. Table 5.

Two distinct cloning strategies were employed based on the specific 5′ UTR architecture. The sequences for ^100^NUS1, ^200^NUS1, ^400^NUS1, ^300^NUS2, and ^300^NUS2-SL were amplified via PCR using primers designed to introduce flanking PstI and XmaI restriction sites (Suppl. Table 6). Following amplification, amplicons were resolved by agarose gel electrophoresis and recovered using gel purification kit (Promega). Purified inserts and the destination vector were digested with PstI-HF (NEB) and XmaI (NEB) and subsequently ligated using T4 DNA ligase. The ^50^NUS1 and ^300^NUS1 constructs were generated via Gibson Assembly into the pSP64 Poly(A) backbone. Primers utilized for these assemblies, incorporating the necessary overlapping homology arms, are detailed in the Suppl. Table 7.

### In Vitro Transcription of GFP Reporter mRNA

Plasmids were linearized downstream of the sequence encoding an A30 poly(A) tail site using EcoRI-HF (NEB). Successful linearization was confirmed via 1% (w/v) agarose gel electrophoresis in TAE buffer. Linearized templates were purified by phenol/chloroform extraction and ethanol precipitation, then resuspended in 50 μL of nuclease-free water (ddH2O).

Transcription reactions were performed using 3.4 µM of in-house purified T7 RNA polymerase and 20-40 pM linear DNA template in the buffer containing 80 mM HEPES pH 7.4, 2 mM spermidine, 40 mM DTT, 24 mM MgCl_2_, 0.4 U/µl murine RNAse inhibitor (NEB), and 7.5 mM each of ATP, GTP, UTP, and CTP.

After 4 hours of incubation at 37°C, RNA was precipitated overnight at-20°C by adding 2.5 M LiCl and 19 mM EDTA. The RNA was pelleted by centrifugation (13,300 rpm, 30 minutes, 4°C), washed with 70% ethanol, air-dried, and resuspended in 50 µL ddH20. mRNA was then desalted twice using 1 mL of G-25 Sephadex® G-25 Medium (Cytiva) equilibrated with ddH2O overnight, centrifuged in spin columns (Bio-Rad).

### Enzymatic m^7^G Capping

For post-transcriptional capping, 10-20 µg of mRNA was treated with Faustovirus Capping Enzyme (NEB) according to the manufacturer’s instructions. Capped products were recovered via precipitation with 2.5 M LiCl and 19 mM EDTA and further purified using the RNA Clean & Concentrator-25 kit (Zymo Research).

mRNA integrity was assessed by electrophoresis on a 1% (w/v) agarose gel in BPTE buffer (100 mM PIPES, 300 mM Bis-Tris and 10 mM EDTA) at 80V for 1 hour. Before loading, samples were denatured by incubation with glyoxal loading dye (Invitrogen) in a 1:1 ratio (v/v) at 50°C for 30 minutes, followed by snap-cooling on ice.

### Cell-Free In Vitro Translation Assays

Two cell-free systems were utilized to evaluate GFP expression.

### Wheat Germ Extract (WGE)

GFP mRNA (160 fmol) was pre-incubated in 50% (v/v) micrococcal nuclease-treated WGE (Promega) with 50 mM potassium acetate and 0.8 U/µL RNAse inhibitor (Invitrogen) for 10 minutes at 25°C. Translation was initiated by adding an amino acid mixture (final conc. 0.08 mM each). All reactions (20 uL final volume) were transferred to 384-well black plates (Corning). GFP fluorescence (λex = 485 nm; λem = 510 nm) was monitored every 128 seconds for 2 hours at 25°C using Tecan Spark High performance Multi-Mode Microplate Reader.

### HEK293T Lysate (ECE-CFPS)

Reactions were prepared with 1.60 pmol GFP mRNA in a buffer containing 50% (v/v) CFPS lysate, 10 mM HEPES pH 7.4, 2 mM DTT, 0.26 mM spermidine, 8 mM amino acid mixture, 10 µg/ml creatine kinase, 21 mM creatine phosphate, 0.2 mM GTP, 0.9 mM ATP, 2 mM Mg(OAc)_2_ and 50 mM KOAc. All reactions (20 uL final volume) were transferred to 384-well black plates (Corning). GFP fluorescence was monitored every 128 seconds for 2 hours at 30°C using Tecan Spark High performance Multi-Mode Microplate Reader.

### Analysis of RNA stability by Real-time quantitative PCR (RT-qPCR)

Translation reactions were halted at the indicated time points (t=0 and 1 hr for WGE; t=0 and 2 hr for HEK293T). Total RNA was isolated via phenol/chloroform extraction followed by ethanol precipitation. The resulting pellets were washed with 70% ethanol, air-dried for 1 hour and resuspended in 50 µl ddH2O. Final RNA purification was performed using RNA Clean & Concentrator-25 kit (Zymo Research). cDNA was synthesized using SuperScript III reagents (Invitrogen) with random primers as per the manufacturer’s protocol. Quantitative PCR (qPCR) was performed using Fast SYBR green PCR master mix, 7500 Fast real-time PCR system sequence detection, and 7500 software version 2.0.1 (Applied Biosystems) following the manufacturer’s instructions. GFP transcripts were quantified with primer pair: 5′ ATGGCCCTGTCCTTTTACCAGA 3’ and 5′ AATGGTTGTCTGGTAAAAGGACAGGGCCAT 3′. Quantitation was performed using the ΔΔCT (threshold cycle) method. Changes in mRNA levels after 1-(HEK293T cell extract) or 2-hour (WGE) translation were quantified relative to amounts of mRNA extracted from cell lysate before translation as 2^-ΔCt^ from respective Ct (cycle threshold) values, where ΔCt = Ct_after_translation_ – Ct_before_translation_.

### Mammalian cell line culture

HEK293T cells were cultured in DMEM media (Gibco) containing 4.5 g/L glucose, and supplemented with 10% FBS (Corning), 100 units/ml Penicillin (Gibco), 100 µg/ml Streptomycin (Gibco), and 2 mM L-Glutamine (Gibco). Cells were grown in a humidified tissue culture incubator at 37°C and 5% carbon dioxide.

### Generation of cells stably expressing GADD34Δ and K3L proteins

A stable cell line endogenously expressing GADD34Δ and K3L was generated as previously described^42^. Wild-type HEK293T cells were co-transfected with the pCMV(CAT)T7-SB100 encoding SB100X (Sleeping Beauty) transposase and the pSBtet-Hygro-GADD34_Δ1-240/P2A/K3L plasmid encoding GADD34 (regulatory subunit of protein phosphatase 1) and vaccinia virus K3L proteins. Transfections were done using Neon transfection system (Invitrogen). pCMV(CAT)T7-SB100 was a gift from Zsuzsanna Izsvak (Addgene plasmid # 34879; http://n2t.net/addgene:34879; RRID:Addgene_34879)^62^. pSBtet-Hygro-GADD34_Δ1-240/P2A/K3L was a gift from Jamie Cate (Addgene plasmid # 196136; http://n2t.net/addgene:196136; RRID:Addgene_196136)^42^. Two days after transfection cells were selected with 300 µg/ml hygromycin (Sigma Aldrich). The stable cell line was regularly maintained with 150 µg/ml hygromycin in growth media.

### Preparation of cell extracts for in vitro translation

4-4.5 million HEK293T cells were seeded in 150 mm plate in DMEM containing 150 µg/ml hygromycin. On the following day 1 µg/ml of doxycycline (Thermo Scientific) was added to the media to induce expression of GADD34Δ and K3L. 24 hours later cells were collected by scraping on ice, washed with ice-cold PBS (Gibco), and resuspended in an equal volume of hypotonic lysis buffer (10 mM HEPES pH 7.4, 10 mM KOAc, 0.5 mM Mg(OAc)2, and 5 mM DTT). Cells were incubated on ice for 45 minutes and homogenized by pushing through a 1-ml syringe with a 27G needle 10 times. Next, the lysate was centrifuged at 15000g for 5 min, and the supernatant was aliquoted and flash-frozen in liquid nitrogen.

## ACKNOWLEDGEMENTS

This work was supported by grants of the National Institutes of Health R35GM141812 (to D.N.E.) and R35GM145283 (to D.H.M.). T.O. was supported by NIH T32GM135134. We thank Regina Cencic, Francis Robert, and Jerry Pelletier for providing hippuristanol.

## AUTHORS CONTRIBUTIONS

T.O., D.H.M. and D.N.E. designed experiments. T.O., O.C. and A.B. performed experiments. T.O. and D.N.E. wrote the manuscript. T.O., O.C., D.H.M and D.N.E revised the manuscript.

## CONFLICT OF INTEREST

Authors declare no conflict of interest.

## Notes

### Competing Interest Statement

The authors have declared no competing interest.

